# Intracranial characterization of gamma in humans in response to auditory gratings

**DOI:** 10.64898/2026.10.04.756534

**Authors:** Divya Gulati, Malla Bhaskara Rao, Ravindranadh Chowdary Mundlamuri, Arivazhagan Arimappamagan, Nishanth Sadashiva, Sanjib Sinha, Raghavendra Kenchaiah, Ajay Asranna, Lakshminarayanapuram Gopal Viswanathan, Supratim Ray

## Abstract

Activity in the gamma frequency range (>30Hz) serves as a neural hallmark of stimulus encoding across multiple sensory modalities. Based on its distinct properties, this activity could be classified as broadband or narrowband oscillations. Visual stimuli, such as a grating, induce narrowband oscillations, which have shown promise as diagnostic markers for neurological disorders. However, such tasks are discomforting for the subjects. These can be made subject-friendly by swapping the stimulus for an auditory one; however, most auditory stimuli elicit a broadband activity. In search of an ideal candidate to induce auditory narrowband oscillations, we previously recorded responses to auditory gratings in electroencephalography (EEG) recordings (Gulati & Ray, 2024) using a passive listening paradigm. We observed a focal sustained broadband increase in high gamma activity (70-150Hz), which was not selective towards any stimulus attributes. Nevertheless, since EEG represents activity of a large population, it remained unclear whether observed activity represented a broadband increase or comprised multiple narrowband rhythms, and whether it was indeed untuned. To elucidate this, we recorded invasively from the regions on and around the superior temporal gyrus, closer to Heschl’s gyrus of seven patients undergoing epilepsy resection surgery. Patients were either under anesthesia or fully awake while we presented auditory gratings through loudspeakers. We found a transient increase in broadband gamma activity (30-100Hz) followed by a sustained suppression of beta rhythm (14-28Hz). This broadband gamma was selective for slower spectro-temporal modulations. Our result compares the responses obtained across different recording scales and corroborates that auditory gratings produce broadband but not narrowband gamma activity.

## Introduction

Sensory-driven cortical processing is typically associated with changes in the gamma band (≥30 Hz) which are observed across various tasks, cortical regions, and species (Adrian, 1942; Canolty et al., 2006; Crone et al., 2001; Gray & Singer, 1989; Miller et al., 2014; Murty et al., 2018; Muthukumaraswamy & Singh, 2013). This activity could be either non-oscillatory or oscillatory in nature, known as broadband or narrowband gamma. In invasive microelectrode recordings, broadband gamma is tightly correlated with the average firing rate of the neurons around the microelectrode (Ray & Maunsell, 2011) and is present for all kinds of visual and auditory stimuli (Bartoli et al., 2019; Crone et al., 2001). It occurs in a frequency range from 30-200Hz without any clear bump in the spectra (Jia et al., 2011). On the contrary, periodic narrowband gamma oscillation appears as a bump in spectra, occurring between the frequency range of 30-80Hz, and is highly dependent on stimulus features (Jia et al., 2011; Ray & Maunsell, 2011). It is predominantly reported in the visual cortex, contingent on presenting specific stimuli such as gratings and hue patches (Murty et al., 2018; Shirhatti & Ray, 2018).

These oscillations are reflective of push-pull interactions between excitatory and inhibitory neurons (Buzsáki & Wang, 2012; Whittington et al., 1995), and when these interactions are imbalanced, as in the case of Alzheimer’s and schizophrenia patients, this visual stimulus-induced gamma becomes weaker (Grent-‘t-Jong et al., 2016; Murty et al., 2021). Thus, it can serve as a valuable tool for diagnosing such disorders. However, such studies involve cumbersome visual tasks, often causing discomfort to subjects (Wilkins et al., 1984). Henceforth, substituting a visual stimulus with an auditory one may make these tasks more manageable. But various stimuli such as pure tones, clicks, noise, frequency sweeps, words, and speech all elicit an increase in broadband activity (Billig et al., 2019; Brugge et al., 2009; Canolty et al., 2007; Crone et al., 2001; Edwards et al., 2005; Griffiths et al., 2010; Lenz et al., 2008; Sedley et al., 2012; Trautner et al., 2006), and thus auditory stimulus-induced narrowband gamma in humans remains elusive.

We hypothesized that a direct analog of visual gratings, auditory ripple stimuli, or auditory gratings (Shamma, 2001) might be an optimal choice to induce narrowband oscillations. These complex broadband sinusoids can be parametrically modulated spectrally and temporally. Owing to the broadband nature of these stimuli, they drive the auditory neurons robustly (deCharms et al., 1998) and have proven to be extensively valuable for characterizing auditory spectro-temporal receptive fields (Kowalski et al., 1996; Massoudi et al., 2015).

Our previous study (Gulati & Ray, 2024) investigated whether these stimuli could induce a narrowband gamma rhythm in electroencephalography (EEG) recordings. We observed that ripples elicited a sustained broadband high gamma response (70-150Hz) across different stimuli, which was non-selective towards any stimulus feature. However, since EEG represents a volume-conducted signal, it was ambiguous whether the broadband gamma observed was multiple narrowband gamma signatures with different center frequencies or an actual wideband increase. Additionally, the absence of tuning of gamma activity could also be attributed to spatial summation across smaller populations with different tuning preferences. For example, visual stimulus-induced gamma measured with EEG has been shown to have much weaker selectivity than gamma recorded from the local field potentials (LFP) (Murty et al., 2018).

Invasive recordings such as electrocorticography (ECoG) could help overcome these limitations as they offer a better spatial resolution and signal-to-noise ratio than EEG (Mukamel & Fried, 2012), and the selectivity of gamma in ECoG is also found to be similar to that observed in LFP recordings (Dubey & Ray, 2020). In this study, we recorded activity from ECoG or stereo-EEG (sEEG) electrodes implanted near Heschl’s gyrus in patients undergoing epilepsy surgery, in response to auditory ripple stimuli, and characterized the response.

## Methods

### Human subjects

Patients with drug-resistant epilepsy who were undergoing brain surgery for seizure foci resection consented to participate in a brief auditory study during the seizure foci boundary marking step during the surgical procedure. These patients presented no symptoms to suggest auditory dysfunction. In some cases, patients were implanted with electrodes semichronically for a few days to identify potential foci. Electrode type (grid or semichronic stereoEEG electrode), electrode grid size, electrode location, and anesthesia management were based solely at the clinical team’s discretion. Due to the experiment, an additional 5 minutes were added per surgery that otherwise lasted for several hours. All patients were provided a written consent form, briefing them about the study. In case of minors (under 18), assent forms were provided to obtain consent from the parents. The experimental protocol was approved by the Institute Human Ethics Committee (IHEC) of the Indian Institute of Science and the Institute Ethics Committee of NIMHANS. Patients whose grid coverage was insufficient, seizure foci were located near Heschl’s gyrus, or who had a noisy recording session, were removed from further analysis. Further, we applied stringent criteria on the spectral responses for selection (see below), based on which seven out of a total of thirty-seven patients (three males, age ∼27.4 ± 7.18 years (mean±std)) were included for final analysis. Out of these, recordings from six subjects were performed during the surgery, and for one subject, the recording was done pre-operatively.

### Setup and data acquisition

ECoG electrodes were platinum disc-shaped, 4mm in diameter, with 2.3 mm exposure (Ad-Tech Medical Instrument, USA). The size of ECoG grids varied from 20 to 40mm in width and 80mm in length, with contacts ranging from 16 to 32. The inter-electrode distance was 10mm. SEEG electrodes were platinum, 2mm cylindrical contacts with 0.86 mm diameter, with 10 mm spacing between the two contacts (Ad-tech). ECoG electrodes were connected to the Brain Products data acquisition system using a 64-channel electrode input box (EIB64-A). The ground electrode was placed near the subject’s shoulder region. The reference electrode was placed on the scalp near the ear, opposite to the craniotomy side. In case of recording from the patient with a semichronic implant, jumper and adapter cable with 1.5mm DIN connectors were used to split the data stream going to the acquisition system used for seizure detection to Brain Products. This was done to avoid disturbing the data stream used for seizure detection. Raw signals were sampled at 1000Hz, filtered between 0.016Hz and 250Hz using filter-order and fifth-order Butterworth filters, respectively. The signals were digitized at 16-bit with 0.1μV/bit resolution. The markers indicating stimulus onset and offset were recorded using a digital I/O card by National Instruments (USB 6210).

### Experimental setting and auditory paradigm

The auditory protocol was run in operating or Video EEG recording rooms with background noise around 50-55dB. When the protocol was run in the operating room, subjects were under anesthesia (refer to Table 1 for details). The patient with a semichronic implant was instructed to listen to sounds passively while sitting upright with eyes closed, with minimal body movement. Heart rate monitor beeps were turned off during the sound presentation. Auditory stimuli were presented using a multidirectional speaker (Marshall Kilburn II) at 75-80dB SPL. Speakers were placed either below the patient’s head near the operating table or in front of the patient with a semichronic implant. The sound stimuli were generated in MATLAB and were presented using the NIMH MonkeyLogic toolbox in MATLAB. We presented spectrotemporally modulated ripple sounds, which are equivalent to multiple amplitude-modulated sinusoids of different frequencies summed up (Kowalski et al., 1996). We used three ripple frequencies (RF – 0, 0.8, 1.6 cycles/octave) and four ripple velocities (RV – 0, 5, 10, 20 cycles/s or Hz) to generate 12 kinds of ripple stimuli. Each ripple stimulus was repeated 20 times and was presented for 800ms, followed by an intertrial interval (ITI) of 800ms. For one subject, the ITI was 200ms. In this subject, an additional stimulus with no amplitude was played for 800ms, which was used as the baseline period for analysis. Further information on how the sounds were generated can be found in the methods section of our previous study (Gulati & Ray, 2024).

**Table 1.** Subject Details.

| *Subject* | *Age* | *Sex* | *Grid* | *Grid Location* | *Anesthesia Details* |
| --- | --- | --- | --- | --- | --- |
| *No.* | *(years)* |  | *size* |  |  |
| 1 | 14 | F | - | 19 stereo EEG electrodes in the left hemisphere, refer to Table 2 for location | Not under any sedatives – Awake recording (5 days post grid implantation surgery) |
| 2 | 30 | M | 4×8 | Laterally along the right superior temporal gyrus | Induction – Fentanyl- 100µg, Thiopentone- 250mg, Vecuronium- 8mg. Maintenance – Desflurane 5-6% in air/oxygen (50:50) plus fentanyl boluses. |
| 3 | 27 | M | 4×8 | Laterally along the left superior temporal gyrus | Induction – Fentanyl- 200µg, Thiopentone- 300mg, Vecuronium- 8mg. Maintenance – Sevoflurane at 2.5% in air/oxygen. Scalp local anesthesia was also given. |
| 4 | 35 | M | 2×8 | Laterally along the left superior temporal gyrus | Induction – Fentanyl- 150µg, Thiopentone- 350mg, Rocuronium- 70mg. Maintenance – Desflurane – 1 MAC (Minimum Alveolar Concentration), Cisatracurium – 15mg/hr |
| 5 | 34 | F | 2×8 | Laterally along the left superior temporal gyrus | Induction – Thiopentone- 300mg, Vecuronium- 8mg, Morphine- 6mg. Maintenance – Sevoflurane at 1 MAC in air/oxygen |
| 6 | 23 | F | 2×8 | Laterally along the right superior temporal gyrus | Induction – Fentanyl- 100µg, Thiopentone- 250mg, Vecuronium- 8mg. Maintenance – Sevoflurane at 1 MAC in air/oxygen. |
| 7 | 29 | F | 4×8 | Slid onto the left temporal cortex region, craniotomy was done on the occipital region (blind placement) | Induction – Fentanyl- 100µg, Propofol- 100mg, Vecuronium- 10mg. Maintenance – Desflurane 3% in air/oxygen. Dexmed - 20µg/hr and Propofol- 100mg/hr |

### Artefact rejection

We employed a pipeline similar to the one used in our previous study to reject artefacts from the EEG data (Gulati & Ray, 2024). Briefly, we first passed each trial through a high-pass filter with a cutoff of 1.6Hz. We then calculated the root mean squared (RMS) value (between −500ms to 1300ms or −200ms to 1000ms (only for trials where ITI was 200ms)) to apply a fixed threshold (with lower and upper cutoffs of 40μV and 400μV) — any trial with the RMS value outside these bounds was considered noisy. Next, we applied thresholding to the remaining trials on their multi-tapered PSD to select the noisy trials, i.e., those deviating by >6 standard deviations in the frequency range of 0 to 200Hz were rejected. We discarded electrodes with more than 30% of all trials marked as noisy and those with PSD slopes less than zero in the range of 56-84Hz. Using this criterion, we rejected 7.44±7.22% (mean±std) electrodes and 5.53±2.48% trials from each electrode per subject.

### ECoG data analysis

We rereferenced data to average referencing for all analyses to remove any common noise. All the data was analyzed using custom-written codes in MATLAB (The MathWorks, Inc., RRID: SCR_001622). We did the spectral analysis using the Chronux toolbox (Bokil et al., 2010, RRID: SCR_005547). For our final analysis, we only used the electrodes that were anatomically closest to Heschl’s gyrus or were in direct vicinity of the ones indicated. We computed power spectral density and time-frequency spectrograms using the multitaper method with a single taper. For time-frequency spectra, we used a moving window of size 250ms with a step size of 25ms, yielding a 4Hz resolution. We calculated the change in power in beta (12-28Hz) and broadband gamma ranges (30-100Hz; 70-150Hz) using the following equation:

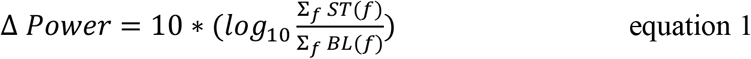

where *ST* and *BL* represent the trial-averaged power during the stimulus and baseline period, respectively. *f* represents the frequencies of interest. For averaging power across frequency bins, we first summed up the power across the frequency range and then converted it to log. The stimulus period for the broadband gamma bands was 50ms to 350ms, and for the beta band was 450 to 750ms. The baseline period was considered between −400ms to −100ms.

We calculated the coefficient of variation (CV) of power during the stimulus period across different stimulus conditions. For this, we calculated a ratio of standard deviation by the average power in the broadband gamma bands for the different kinds of stimulus conditions for each selected electrode. CV values for the EEG data were taken from Gulati & Ray (2024).

### Statistical analysis

We compared the CV values across ECoG electrodes with those obtained for EEG data across subjects using the Wilcoxon rank-sum test with the null hypothesis stating medians came from the same population.

### Data and code availability

Spectral analyses were done using the Chronux toolbox (version 2.10) available at http://chronux.org. Data will be made available to the reader on reasonable request.

## Results

We recorded intracranial electrocorticography data from seven patients while the subdural grid or the sEEG electrodes were placed or implanted near Heschl’s gyrus along the superior temporal gyrus. While recording this data, we played auditory grating stimuli while the subjects were under general anesthesia, except one subject, for whom recordings were done while the subject was awake and had eyes closed. We presented 12 different types of auditory grating stimuli, each presented for 800ms.

### Ripple stimuli induce broadband gamma

Figure 1 shows the trial-averaged event-related potential (ERP) responses and time-frequency spectrograms in response to auditory grating stimuli for all the recorded electrodes in one example subject, who passively listened to these sounds with eyes closed. We recorded data from four sEEG electrodes with varying numbers of contacts (four to six). Electrodes were implanted semichronically in the temporal and occipital cortex; the exact locations of the electrodes are mentioned in Table 2. Prominent ERP responses were obtained at the electrodes located near the posterior superior temporal gyrus. The strongest N1 wave peak, a negative deflection around ∼90ms (Sur & Sinha, 2009), was obtained for the electrode located at the junction of the superior temporal gyrus and Heschl’s gyrus. For the same electrodes, we observed a transient increase in broadband gamma activity (30-100Hz) followed by a sustained suppression of beta oscillations (12-28Hz). The increase in broadband gamma lasted for ∼250ms post stimulus onset, whereas beta suppression lasted until stimulus offset. Importantly, there were no narrowband oscillations in the gamma range, which are represented as horizontal red bands in the time-frequency plots as observed in the visual cortex (see Figure 1 of Gulati & Ray, 2024). Therefore, unlike visual gratings, these auditory gratings do not produce narrowband gamma oscillations but only a broadband gamma response followed by low-frequency suppression.

**Figure 1.**
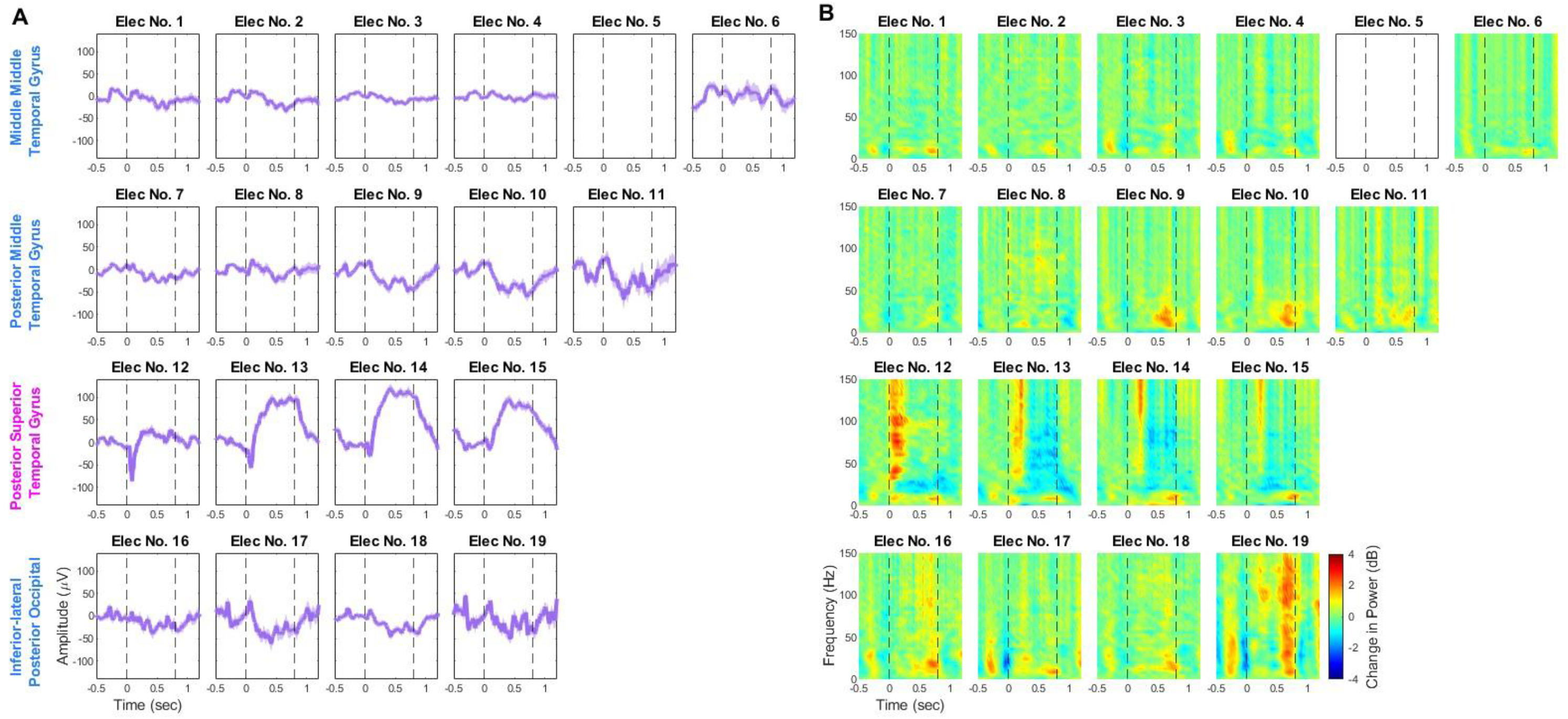
Event-related potential (ERP) and time-frequency change in power spectrograms for different electrodes implanted in a representative subject. The power spectra are averaged across different stimulus conditions. (A) Grand averaged ERP waveforms for 4-contact sEEG electrodes implanted in four brain regions are shown across the rows (Middle middle temporal gyrus, Posterior middle temporal gyrus, Posterior supra-temporal gyrus, Inferior-lateral posterior occipital region). (B) Time-frequency spectrograms for the same. The colorbar represents the log power ratio in decibels. The electrode with an empty plot was rejected from the analysis.

**Table 2.**
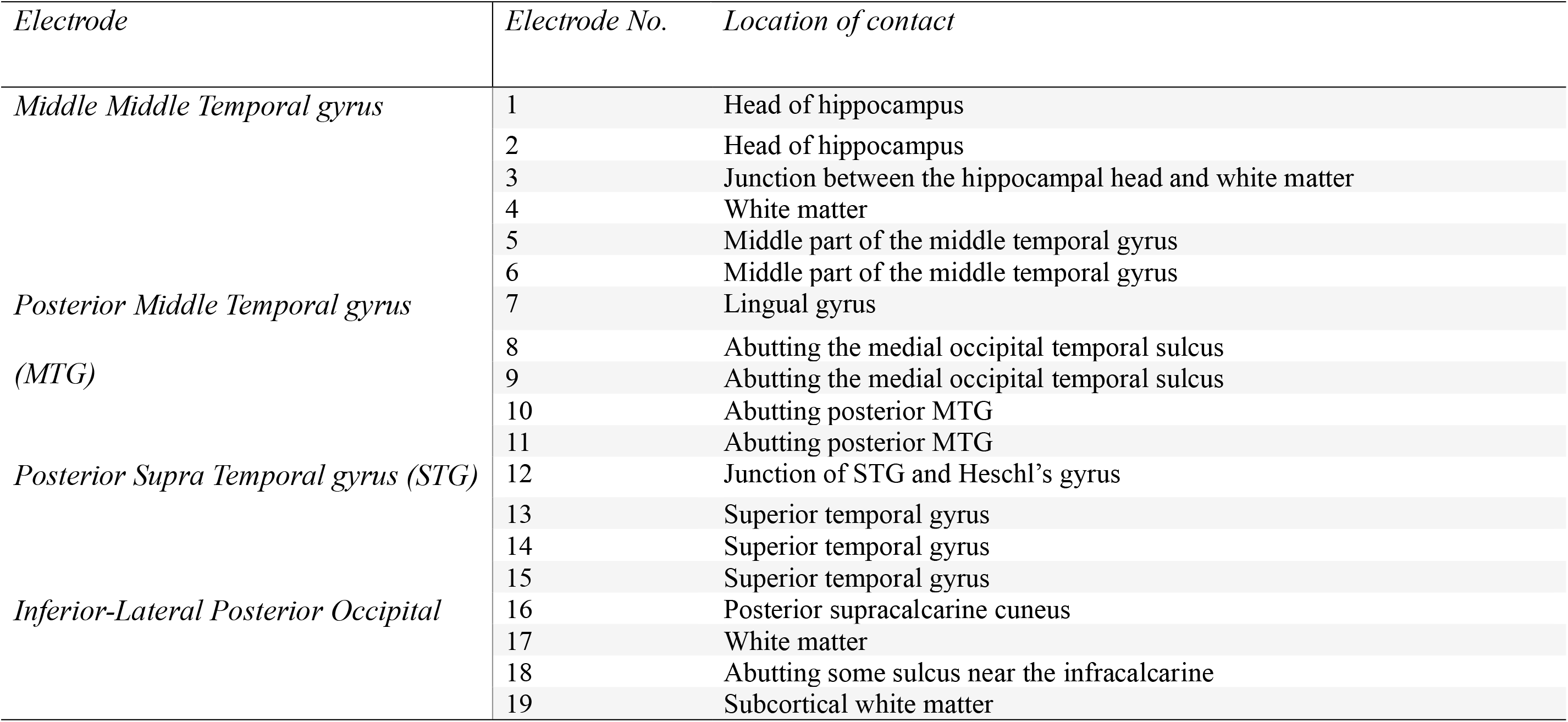
Showing locations of each contact for the four sEEG electrodes used for the representative subject.

Figure 2A-B shows the average change in power in gamma and beta frequencies across the electrode grids implanted for all seven subjects. We selected only those electrodes for analysis (dashed black boxes in Figure 2A-B) that were lying closest to Heschl’s gyrus (yellow boxes) or were in the direct vicinity of these electrodes and additionally also showed both the increase in gamma, followed by a decrease in beta oscillations in their respective time-frequency spectrograms. Except for one subject, for whom the grid placement on the cortex was blind, as craniotomy was done on the occipital region and the grid was slid over the temporal cortex, the electrode was selected based on the presence of similar neural signatures. This resulted in 12 electrodes across seven subjects, marked with dashed black rectangles. Figure 2C shows the time-frequency spectrograms for each of the subjects averaged across the selected electrodes. We observed slight differences in gamma synchronization and beta desynchronization latency across subjects. This could be attributed to minor variations in individual microanatomy and the precise grid location across the subjects. Also, it could be observed that these responses are similar irrespective of the state of arousal or the type of anesthetic (shown in Table 1) that was administered to the subject. These results match those obtained in EEG using the same stimuli (Gulati & Ray, 2024), with the only difference being that we obtained a sustained broadband activity for the entire stimulus instead of the transient response observed here. None of these subjects showed a narrowband gamma response.

**Figure 2.**
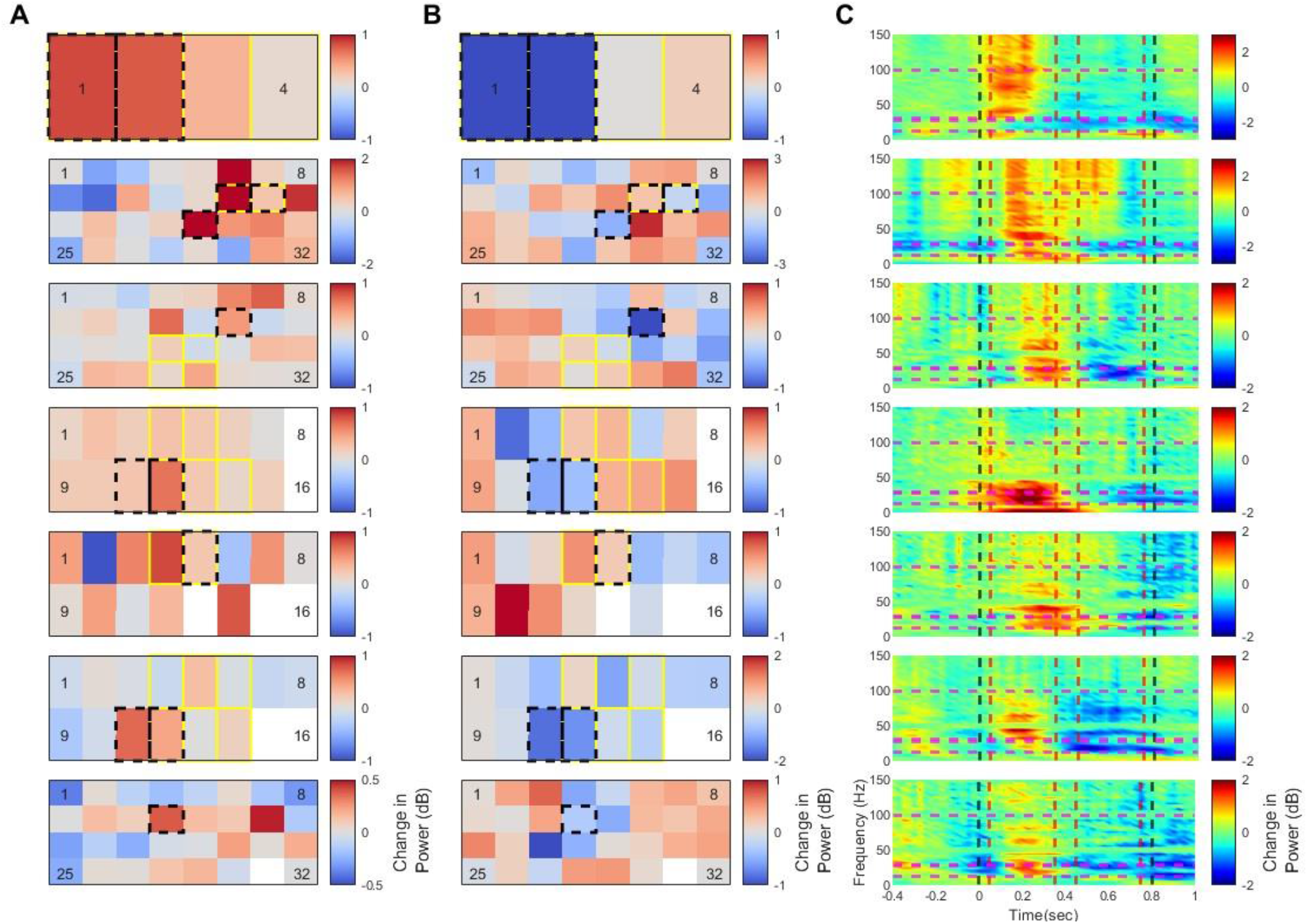
Change in power for stimulus conditions compared to baseline condition (−400 – 0ms), pooled across all stimulus conditions for each subject. (A) Heatmaps showing mean change in power in the broadband gamma band (30-100Hz) during the stimulus period (50-350ms) for each electrode of the sEEG electrode or ECoG grid for each subject. Each row represents one subject. The colorbar represents frequency and time-averaged change in power in dB. Electrodes marked in yellow and black represent the electrodes indicated by the surgeons as closest to Heschl’s gyrus and the selected electrodes for the analysis, respectively. Electrodes filled with white colour were the electrodes labelled as noisy. (B) Heatmaps showing the mean change in power in the beta band (12-28 Hz) during the stimulus period (450-750ms). The rest is the same as in panel A. (C) Time frequency spectrograms averaged across the selected electrodes for each participant. The dashed vertical black lines indicate stimulus onset and offset. The dashed horizontal magenta and vertical red lines mark the frequency and time bins used for averaging power in gamma and beta bands in panels A and B, respectively. The colorbar represents the log power ratio in decibels.

### Stimulus selectivity of broadband gamma could be observed in ECoG

Previous invasive studies in the visual domain have found gamma to be stimulus-selective (Jia et al., 2011; Murty et al., 2018), but such selectivity is often diminished in non-invasive EEG recordings (Gulati & Ray, 2024; Murty et al., 2018). This is attributed to the relatively poor spatial resolution and volume conduction effects in EEG recordings. The lack of stimulus selectivity of the gamma response in EEG has also been found for the auditory grating stimuli (Gulati & Ray, 2024). So, we tested with invasive electrocorticography recordings whether we could find broadband gamma to be selective for any particular feature of auditory grating stimuli. We plotted the change in power time-frequency spectrograms for different kinds of auditory grating stimuli for one example subject (Figure 3), averaged across the selected electrodes. For this subject, we observed the strongest gamma responses for auditory gratings with a ripple frequency of 0.8 cycles/octave and ripple velocity of 5Hz. We observed similar responses across all subjects. Figure 4A shows the time-frequency spectrograms averaged across the 12 final set of electrodes obtained across the 7 subjects. To quantify the selectivity, we calculated the coefficient of variation for gamma (30-100Hz) power during the stimulus period (50-350ms) for each electrode (Figure 4B). We obtained CV values for broadband gamma in intracranial recordings (0.18±0.097 - mean±std) larger than those obtained for the broadband gamma power (70-150Hz in the time range of 250-750ms; CV value: 0.066 ± 0.004 – mean±std) in EEG recordings (Gulati & Ray, 2024). Since in the EEG data CV values were calculated for a frequency range between 70-150Hz, we calculated CV values for the same range as well. These were also found to be larger in our data (0.196±0.1004-mean±std). To statistically validate whether the CV values are significantly different than those found in EEG recordings, we performed a Wilcoxon rank-sum test to compare the medians, which was found to be significant for both broadband gamma ranges (30-100Hz: p = 4.8024e-05 and 70-150Hz: p = 1.295e-06), indicating that gamma was much more selective in intracranial recordings than in EEG. To check which stimulus features are preferred by activity in the gamma band, we noted the stimulus condition for which maximum gamma power (30-100Hz) was observed across all the selected electrodes (Figure 4C). We observed that electrodes overall preferred the stationary auditory grating stimuli with ripple velocity of 0Hz and preferred ripple frequencies of 0.8-1.6cycles/octave. The selectivity of gamma matches the tuning observed for ripple stimuli in ECoG, fMRI, and psychoacoustic studies (Chi et al., 1999; Hullett et al., 2016; Langers et al., 2003; Schönwiesner & Zatorre, 2009; van der Willigen et al., 2024).

**Figure 3.**
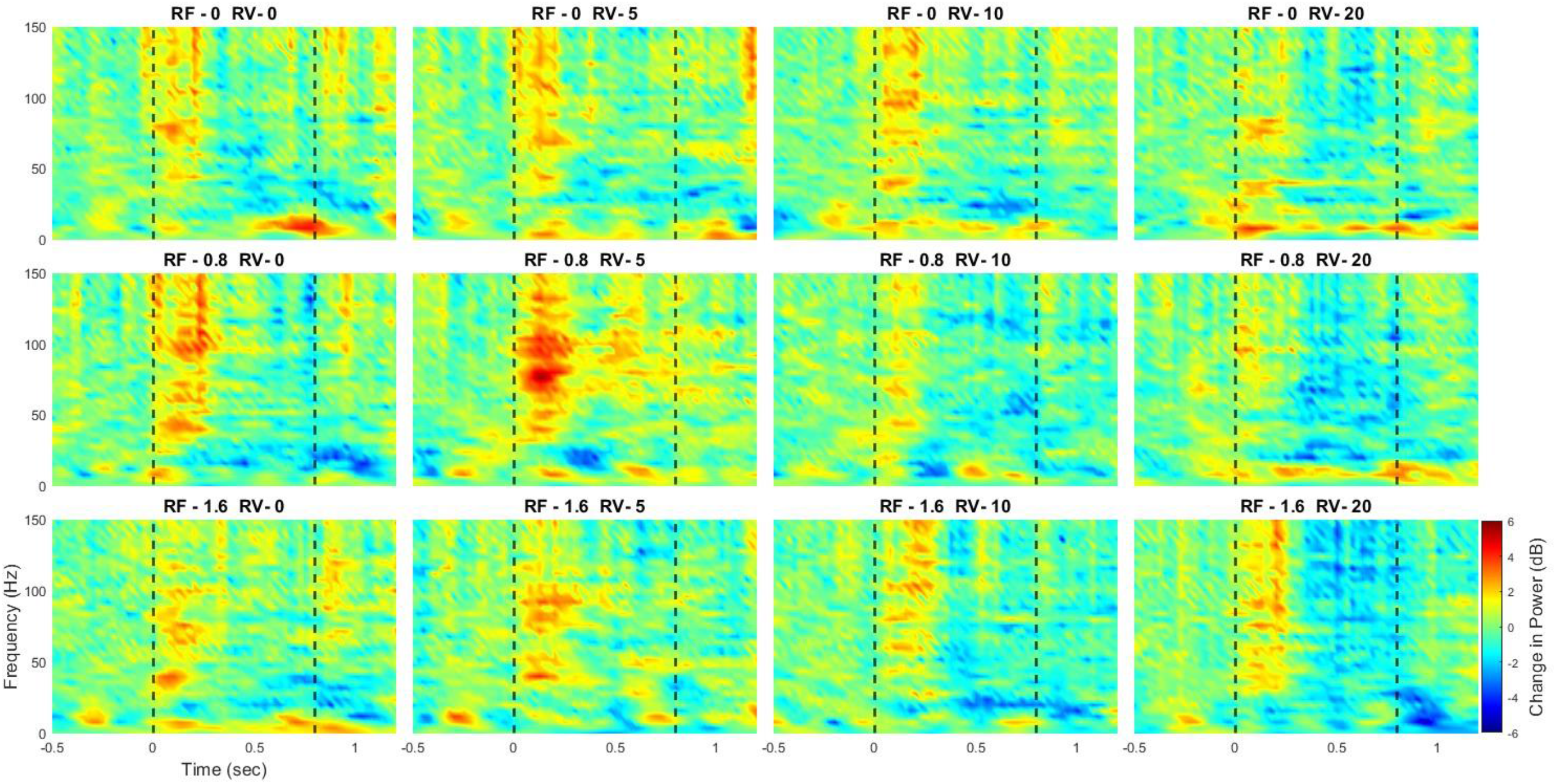
Time-frequency change in power spectrograms for the representative subject under different ripple stimulus conditions. The power spectra are averaged across the selected electrodes (N=2) for the subject. Across the row, stimulus ripple frequency is increasing, and ripple velocity is increasing across the column. The dashed vertical lines (black) indicate the stimulus onset and offset (0-800ms). The colorbar represents the log power ratio in decibels.

**Figure 4.**
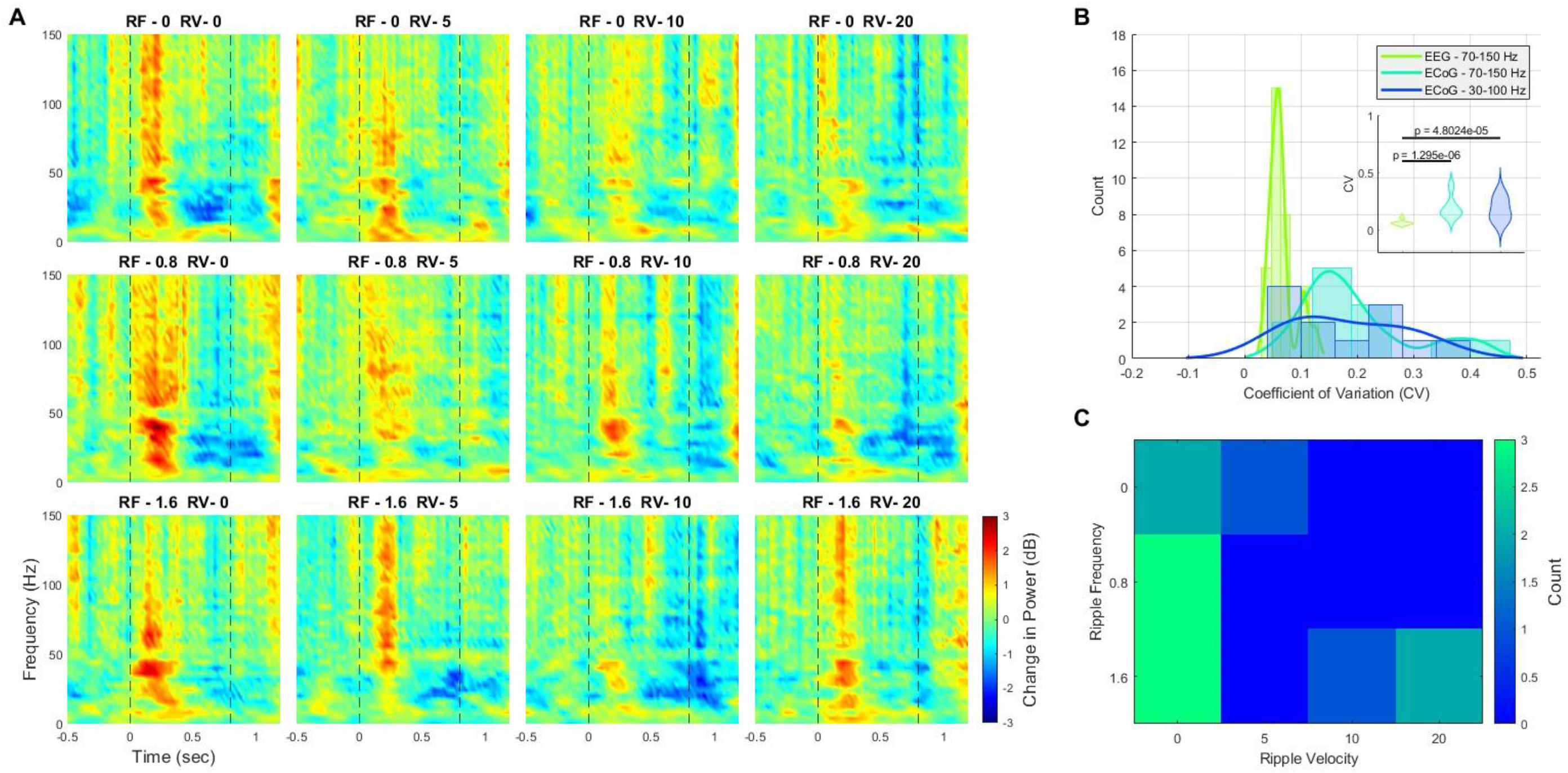
Stimulus selectivity of broadband gamma activity in EcoG. (A) Time-frequency change in power spectrograms for the different ripple stimulus conditions averaged across the selected electrodes. The panel arrangement is the same as in Figure 3. (B) Histogram showing coefficient of variation (CV) values calculated for power in two frequency ranges (30-100Hz and 70-150Hz; time-period – 50-350ms) for each selected ECoG electrode. A histogram showing CV values for each subject averaged across all EEG electrodes for power in the frequency range of 70-150Hz during a 250-750ms time duration is also shown in green. Statistical significance was tested using the Wilcoxon rank-sum test between ECoG and EEG CV value distribution. (C) The heat map shows the number of electrodes that have maximum power during the stimulus period (frequency range: 30-100Hz; time-period: 50-350ms) for that particular ripple condition.

## Discussion

We played auditory grating stimuli to patients either under anesthesia or while they were awake, and recorded either ECoG or sEEG (stereo EEG) signals from the temporal cortex near the sylvian fissure. We did not observe any sustained narrowband gamma rhythm, but instead observed a transient broadband gamma increase that was selective for stimulus features, preferring slowly modulating ripple stimuli in both spectral and temporal axes. This tuning was significantly higher in intracranial recordings compared to comparable recordings using EEG, as done in a previous study (Gulati and Ray, 2024). Along with gamma, these stimuli elicited a sustained beta suppression.

### Comparison with previous invasive studies

In microelectrode studies, broadband gamma has been shown to be tightly correlated with the average firing rate of neurons around the microelectrode (Ray et al., 2008; Ray & Maunsell, 2011). This does not necessarily imply that it is due to spike-bleedthrough alone, since synaptic activity that leads to spiking is also likely to contribute to the broadband gamma (Ray & Maunsell, 2015), and it is difficult to dissociate between the synaptic and spiking contributions in extracellular recordings (Reimann et al., 2013). Indeed, some studies have shown that an increase in broadband activity serves as a proxy of robust local cortical activation generally reflective of asynchronous dendritic inputs (Guyon et al., 2021). Generators of broadband activity are found to be in supragranular layers (Leszczyński et al., 2020); therefore, this activity is evident in ECoG and EEG recordings. At the level of ECoG/EEG, broadband gamma could reflect more synchronous activity since the contribution of synchronous events increases with an increase in the population size that contributes to the signal (Ray et al., 2008). Several previous studies involving presentation of stimuli such as phonemes, words, syllables, and sentences have reported a similar transient increase in broadband gamma activity followed by a sustained suppression in alpha and beta bands (Billig et al., 2019; Canolty et al., 2007; Crone et al., 2001; Edwards et al., 2009; Nourski, Steinschneider, Rhone, Kovach, et al., 2021; Nourski et al., 2022).

This increase and decrease in activity across different bands could also reflect feedforward and feedback processing of complex stimuli (Billig et al., 2019; Nourski et al., 2022). This is consistent with some predictive coding studies, which suggest that feedback related to the prediction is mediated through low-frequency alpha and beta rhythms, while feedforward signals representing prediction errors are mediated using gamma rhythms (Bastos et al., 2020). The desynchronization in the lower frequency range has also been reported to get stronger with increasing task engagement (Nourski et al., 2022), indicating that lower frequencies might be involved in reducing task-irrelevant processing. A previous study (Hullett et al., 2016) that presented moving ripple stimuli also observed a transient increase in high-gamma response (70-150Hz) in the supratemporal gyrus, with no analysis reported for other frequency bands.

### Comparison to EEG results

We observed a transient broadband increase instead of a sustained increase for the entire stimulus duration, as previously observed with EEG (Gulati & Ray, 2024). This difference could arise due to the various sources contributing to the ECoG and EEG measurements. A previous study using speech stimuli showed a transient broadband increase in the primary auditory cortex, whereas auditory association areas show a sustained broadband increase (Giraud & Poeppel, 2012). Another study, which looked at responses to speech stimuli along the length of Heschl’s gyrus, observed sustained high gamma broadband activity (70-150Hz) in the posteromedial region of Heschl’s gyrus, which is buried deep within the gyrus. Conversely, transient high gamma increase and sustained suppression in alpha and beta ranges were observed in electrodes in the anterolateral region of Heschl’s gyrus, which is closer to the cortical surface (Billig et al., 2019). As EEG represents a summed up volume-conducted signal (Nunez & Srinivasan, 2006), the sustained response might also be built from the gamma responses originating from multiple auditory regions at different latencies. Previous studies where sEEG recordings were made from multiple auditory regions had shown that the onset latency of broadband gamma varies across regions (Nourski et al., 2014, 2022).

Unlike in EEG, where we observed an increase in the 70-150Hz frequency range, with intracranial recordings, we observed an increase starting from 30Hz. These results align with those previously observed in response to word stimuli, where a reduction in peak frequency of gamma was noted when subjects were awake versus anesthetized (Krom et al., 2020). The same study also noted an increase in gamma power and a reduction in alpha/beta desynchronization in the anesthetized state compared to the awake state. However, we could not compare power changes since we do not have awake and anesthetized invasive recordings or the EEG recordings from the same subjects. Therefore, further investigations are needed to understand these diverse responses.

### Stimulus Selectivity

This tuning of broadband gamma matched that observed in previous fMRI studies showing stronger voxel activation towards certain ripple stimuli (Langers et al., 2003; Schönwiesner & Zatorre, 2009). This is probably because gamma power is known to be tightly coupled to the blood oxygenation level-dependent signal in visual and auditory cortices (Magri et al., 2012; Mukamel et al., 2005). The spectrotemporal tuning of gamma has been shown to be organized across the supratemporal gyri, with different regions preferring different stimulus properties (Hullett et al., 2016).

The visual stimulus-induced broadband (as well as narrowband) gamma is selective towards a variety of stimulus features, such as contrast (Bartoli et al., 2019), size (Dubey & Ray, 2020), and drift rate (Salelkar et al., 2018). What all stimulus features auditory broadband gamma is tuned towards needs further examination. Visual stimulus-induced high gamma activity in MEG has been shown to weaken in Schizophrenic patients (Grützner et al., 2013). However, it remains to be seen whether auditory broadband gamma also changes with neurological disorders, and thus, whether it can also be used as a biomarker.

### Effect of Anesthesia

Interactions between glutamatergic and GABAergic local circuits are instrumental in the generation of narrowband gamma oscillations (Cardin et al., 2009; Whittington et al., 1995). Since most anesthetics used in our study, such as desflurane, sevoflurane, and propofol, act on GABA receptors (Franks & Lieb, 1994; Garcia et al., 2010), they can modulate gamma oscillations. Early reports of stimulus-induced gamma oscillations were based on recordings performed under anesthesia (Gray & Singer, 1989) and have since been extensively characterized under anesthesia (Henrie & Shapley, 2005; Jia et al., 2011), indicating that sensory processing remains largely maintained under anesthesia. Gamma burst duration and peak frequency distribution of gamma bursts are reported to remain similar across arousal states (Xing et al., 2012), although anesthesia decreases the peak frequency and also modulates the power of stimulus-induced gamma when compared to that recorded during awake conditions (Krom et al., 2020; Lozano-Soldevilla et al., 2014; Nourski, Steinschneider, Rhone, Krause, et al., 2021; Saxena et al., 2013, 2021; Xing et al., 2012). Additionally, power changes at lower frequencies are also noted (Xiong et al., 2024). During anesthesia, a reduction in spiking activity in the auditory cortex has been noted (Elhilali et al., 2004; Krom et al., 2020). However, the fine structure of the spectro-temporal receptive fields of primary auditory neurons is found to be intact (Elhilali et al., 2004), indicating that the selectivity of neurons should not change with the arousal state of the subject. Since the effects of anesthesia are shown to differ across interneuron subtypes, the tuning of gamma responses towards select features might change.

The responses of somatostatin (SOM) neurons are reported to be significantly suppressed compared to those of parvalbumin neurons (Adesnik et al., 2012). Therefore, tuning of stimulus features that depend on the activity of SOM neurons, such as tuning of neurons towards sound frequency (Kato et al., 2017; Kumar et al., 2019), may change under anesthesia.

We note that an inevitable caveat of our study is that these recordings were made from epileptic patients, so the conclusions may vary for a healthy population. However, we obtained consistent results across all patients, even though they had varying case histories and were under different kinds of anesthetics and other medications during the recording. Further, EEG recordings from healthy subjects revealed a similar broadband response (Gulati and Ray, 2024). Even though the auditory stimuli we used failed to elicit the narrowband gamma response typically observed in visual areas with visual gratings, the distinct pattern of broadband gamma and low-frequency suppression elicited by these stimuli could convey important information about the way the brain processes auditory information.

## Author Contributions

D.G., S.R., and M.B.R. conceived the research idea and designed the experiments; M.B.R., R.C.M., A.A., N.S., S.S., R.K., A.A., and L.G.V. recruited the subjects; D.G. collected the data; D.G. and S.R. analyzed the data; and D.G., M.B.R., R.C.M., and S.R. wrote the paper.

## Acknowledgements

This work was supported by Wellcome Trust/DBT India Alliance Senior fellowship (IA/S/18/2/504003) to S.R., IISc-IoE (Institution of Eminence) grant to S.R., a grant from Pratiksha Trusts to S.R., and the Council of Scientific and Industrial Research (CSIR) senior research fellowship to D.G.. We acknowledge the doctors and nursing staff at NIMHANS for helping with the experimental setup.

